# EMTscore infers divergent EMT pathways from omics data and enables rapid screening for correlated gene sets

**DOI:** 10.64898/2026.01.27.702045

**Authors:** Haimei Wen, Leonidas Bleris, Tian Hong

**Affiliations:** Department of Biological Sciences. The University of Texas at Dallas. Richarson, Texas 75080. USA; Department of Bioengineering. The University of Texas at Dallas. Richarson, Texas 75080. USA

## Abstract

**Summary:** Quantitative analyses of epithelial-mesenchymal transition (EMT) have been widely used in several areas of biomedical sciences due to its importance in development and cancer progression, but its multi-contextual nature requires standardization and implementation of gene set scoring methods beyond capacities of conventional tools. We developed EMTscore, a package that provides an efficient implementation of unbiased scoring methods for multiple EMT pathways using individual single-cell or bulk omics data, and the package allows rapid screening for relationships between EMT and other cellular processes.

**Availability and Implementation:** EMTscore is available from GitHub https://github.com/wenmm/EMTscore under the GNU General Public License, and it will be deposited to Zenodo upon acceptance. It is also under review at Bioconductor.

**Contact:** Tian Hong

## 1. Introduction

Epithelial-mesenchymal transition (EMT) is the cellular process in which epithelial cells with tight cell junction and apical-basal polarity transform into motile mesenchymal cells. EMT is responsible for key steps of development, including gastrulation and organogenesis (Nakaya and Sheng, 2013). EMT is also activated in diseases progressions such as fibrosis and metastasis. In particular, cancer cells can exploit the gene regulatory network of EMT and the invasiveness of the mesenchymal (M) cells. Recent experimental and theoretical studies have shown that EMT involves multiple intermediate states rather than a binary switch (Hong, et al., 2015; Yang, et al., 2020), and partial EMT was shown to be crucial for cancer progression (Pastushenko, et al., 2018). In addition, divergent transcriptional programs of EMT can be activated in fate changes towards multiple cell subtypes/states even in the same disease (Groves, et al., 2022; Youssef, et al., 2024). The importance and complexity of EMT have gained substantial significant attention from multiple subfields of biomedical sciences. Over the past decade, several thousand EMT-related research articles have been published each year (**Figure 1A**, gray and green).

**Figure 1.**
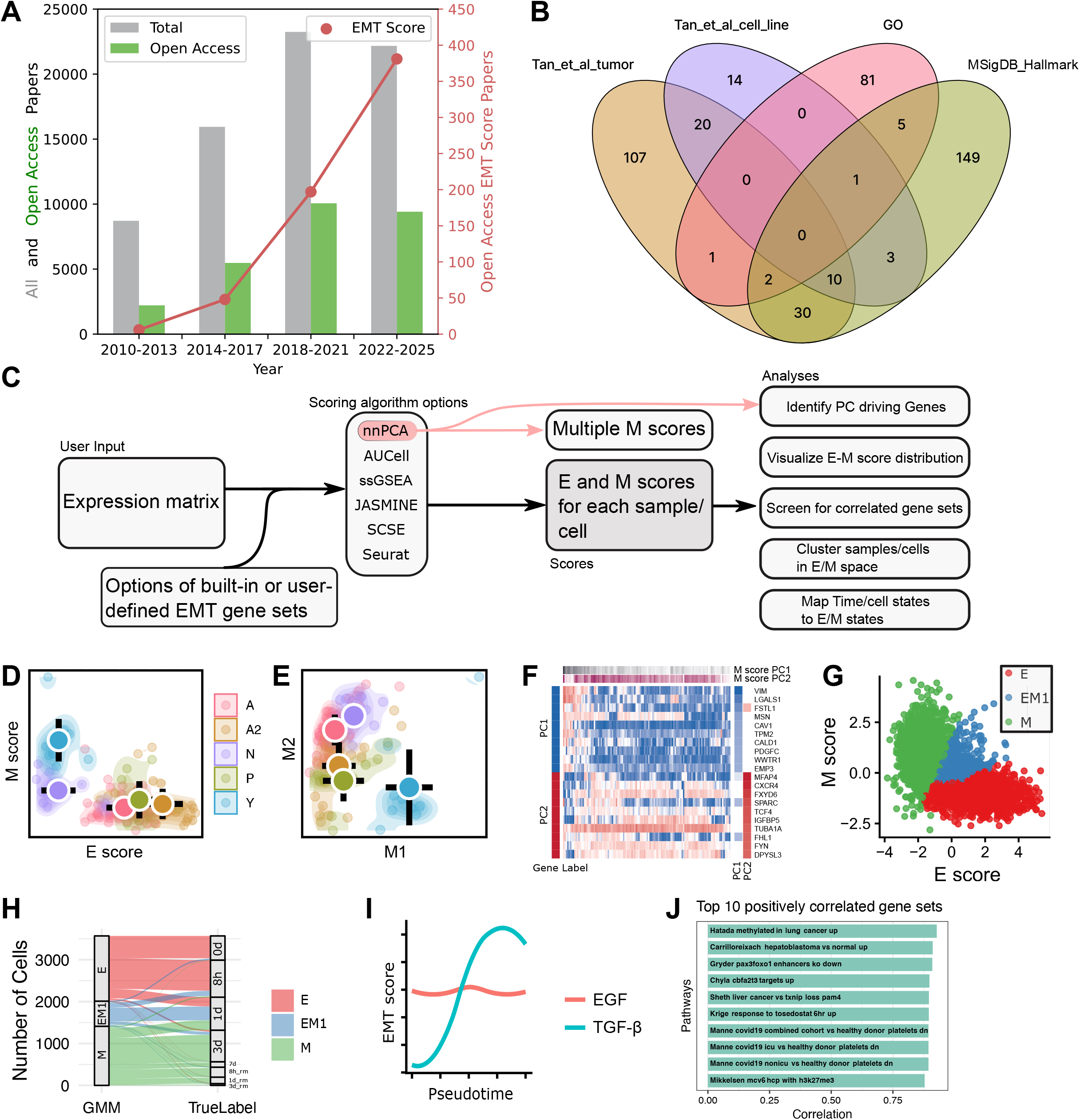
Motivation and overview of EMTscore. **A**. EMT-related publication counts with respect to time. Articles with both keywords “epithelial” and “mesenchymal” mentioned in abstracts were extracted from PubMed (gray) as of December 9, 2025. Open access, original research articles (green) were identified from the total pool. Among open access articles, papers with keyword “EMT score” were further identified (red). **B**. Venn diagram showing the overlaps of four widely used EMT gene sets. **C**. Workflow of EMT score package. **D**. nnPCA-based E and M scores for 120 SCLC bulk RNA-seq samples. **D**. nnPCA-based M1 and M2 scores. **F**. Expression levels of top genes contributing to M1 and M2 scores. **G**. GMM-based clustering of scRNA-seq data for A549 cells treated with TGF-β in E-M space. **H**. Sankey plot showing the mapping of time points and EMT states from GMM clusters. **I**. E and M1 scores as functions of pseudotime for two types of treatments. **J**. Top 10 gene sets whose scores are positively correlated with EMT (M1) scores for TGF-β-treated A549 cells.

Due to the multistate nature of EMT and its diverse physiological contexts, EMT cannot be accurately quantified with a single or a handful of genes (Yang, et al., 2020). Gene sets analysis is therefore widely used to generate “EMT scores” to assess the degree of EMT in different settings (**Figure 1A**, red) (Panchy, et al., 2022; Tan, et al., 2014). Scoring EMT with high-throughput experiments, including single-cell RNA-sequencing (scRNA-seq), enabled both discoveries for fundamental understanding of EMT and development of new therapeutic strategies (Cassier, et al., 2023; Elamin, et al., 2022; Youssef, et al., 2024). Meanwhile, to facilitate characterization of EMT pathways involving complex gene networks, various databases and bioinformatic tools have been developed to date (Vasaikar, et al., 2021; Zhao, et al., 2015; Zhao, et al., 2017). However, these resources do not meet the increasingly complex gene set analyses that researchers are performing. In particular, while general gene set enrichment methods are useful for quantifying EMT progression through scores computed with a defined EMT gene set, the research in this field has been challenged by several key issues. First, different EMT gene sets have been used in different studies (Carbon, et al., 2021; Liberzon, et al., 2011; Panchy, et al., 2022; Tan, et al., 2014), and these gene sets have very limited overlaps (**Figure 1B**), whereas the current gene set analysis tools do not allow easy access to diverse gene sets or fast tests with multiple gene sets. Secondly, discoveries and visualization of divergent activations of subsets of EMT genes require ad hoc methods with which reproducing key results from bioinformatic analysis is difficult (Youssef, et al., 2024). Finally, the mapping from standard cell states annotation or clustering results to biologically meaningful EMT states requires manual inspection, which has limited efficiency and reproducibility.

To overcome these issues, we developed an R package, named EMTscore, that enables easy, simultaneous use of multiple popular EMT gene sets and scoring algorithms, as well as in-depth (single-cell) RNA-seq data analyses of multistate EMT and their visualizations with publication-quality plots. We implemented a method based on nonnegative principal component analysis (nnPCA) that enables automatic detection of groups of divergently activated EMT samples and genes in the same dataset. In addition, the toolbox includes identification of EMT states in low-dimensional functional space and its mapping to user-defined cell types or clusters from standardized methods. Finally, our package can screen gene sets whose expressions are correlated with EMT progression, and this reveals the relationship between EMT and other biological processes in an efficient manner.

## 2. Software Description

### 2.1 Implementation of algorithms for computing EMT scores

To perform scoring of epithelial (E) or mesenchymal (M) gene set activities across transcriptomes (bulk or single-cell samples), EMTscore requires an expression dataset provided as a matrix file, a Seurat object, or a SingleCellExperiment (SCE) object. In addition, EMTscore provides several widely used E/M gene sets which can be replaced by custom gene sets provided by users (**Figure 1C**, left).

EMTscore is mainly designed for complex downstream analyses with EMT scores (described later). To achieve this goal, we implemented multiple scoring algorithms that allow flexibility for choices of popular gene set enrichment methods. In addition, a default scoring algorithm based on nonnegative PCA (nnPCA) enables automated detection of multiple EMT programs (Panchy, et al., 2021; Sigg and Buhmann, 2008) (**Figure 1C**, middle). While this note is not evaluative regarding the algorithmic options, we briefly describe the advantages of the implemented methods here. nnPCA is a fast method based on variance of data points. Its unique advantage of providing multiple axes (e.g. multiple scores) with ranked variances explained makes it suitable for revealing multiple EMT programs in the same dataset. However, nnPCA does not give a statistical test for gene set enrichment even when there are a control and a treatment group in the dataset. In contrast, the widely used single-sample gene set enrichment analysis (ssGSEA) method uses Kolmogorov-Smirnov (KS) test for determining significant enrichment (Barbie, et al., 2009), but the method is inefficient to handle large datasets such as scRNA-seq data. Other more recently developed methods, such as AUCell, SCSE and JASMINE, are either based on gene ranks or normalized sums of expression (Aibar, et al., 2017; Noureen, et al., 2022; Pont, et al., 2019). They are more efficient than ssGSEA but their statistical rigor depends on the context, and they do not allow multiple gene scores from the same dataset. We have implemented each of the methods mentioned above, and the default method is nnPCA which will enable some of the downstream analyses.

EMTscore allows users to choose E and/or M gene sets from commonly used sources (**Figure 1B**). A default set of E/M genes from Panchy et al. can be used. This collection combines a data-derived set of EMT genes and additional, widely recognized transcription factors controlling EMT (Panchy, et al., 2022).

### 2.2 Analysis of multiple EMT transcriptional programs

Upon completion of nnPCA-based scoring, EMTscore extracts multiple leading principal components (PCs) for M scores. Similar to conventional PCA, these PCs are ranked based on their variances explained. For some analyses, one M axis (i.e. the leading PC based on a M gene set) is used, and each sample/cell receives one M score and one E score (**Figure 1D**). To detect divergent EMT (M) programs, two or more PCs from M scores are used. Typically, the first M PC (M1) and the second M PC (M2) can be used to describe the two directions of M programs driving the variance of the data points (**Figure 1E**). Genes whose expressions contribute to M1 and M2 can be extracted based on their rotations (loadings) (**Figure 1F**).

EMTscore visualizes the progression of EMT in the space of E-M for conventional EMT scores, and in the space of M1-M2 for divergent EMT programs. Scatter plots with various options are provided for these visualizations (**Figure 1D** and **E**).

### 2.3 Detection of EMT states with Gaussian Mixture Model

In RNA-seq data, samples are sometimes with experimental labels (e.g. time points). In addition, for scRNA-seq datasets that typically contain a large amount of samples, clustering information is readily available from commonly used packages such as Seurat (Stuart, et al., 2019). However, the relationship between these cluster labels and EMT states is often unclear. To provide insights into this question, EMTscore uses Gaussian Mixture Model (GMM) to cluster cells in E-M space in which the clustering information is directly interpretable biologically. Next, the package detects the extreme E and M subpopulations as well as hybrid EMT subpopulations relative to the dataset (**Figure 1G**). A mapping of the samples’ original labels to EMT states is visualized with a Sankey plot (**Figure 1H**) and the numerical distributions are reported accordingly. In addition, single-cell level labels, such as pseudotime inferred from other packages, can be analyzed in relation to EMT scores (**Figure 1I**).

### 2.4 Screening for cellular processes correlated with EMT

EMT has been shown to have crosstalk with other important cellular processes such as alteration of motility and invasiveness (Jolly and Huang, 2014; Wong, et al., 2014). It is therefore of interest to search for gene set scores that are correlated and/or anticorrelated with EMT scores. Leveraging the high efficiency of scoring methods such as nnPCA, we implemented a screening function for searching through MSigDB (7561 curated gene sets in C2), and ranking gene sets that are highly correlated/anti-correlated with EMT scores (in the case of nnPCA, EMT scores are M1 scores by default but they can be replaced by M2 scores) across samples/cells. EMTscore reports the top positively correlated gene sets and top negatively correlated gene sets separately as lists and bar charts illustrate their Pearson correlation coefficients (**Figure 1J**).

## 3. An example analysis pipeline

We first use an example bulk RNA-seq dataset containing a collection of transcriptomes from 120 small cell lung cancer (SCLC) cell lines with various known tumor subtypes (A2, A, N, P and Y) (Cerami, et al., 2012; Groves, et al., 2022). To investigate the relationship between the subtypes of SCLC and EMT states, we computed E and M scores with each method described above, and we found that A2 was an extremely E-like subtype, whereas Y was extremely M-like (**Figure 1D**). The EMT identities for other subtypes were not clear from a single M score. With nnPCA, we found that there was a divergent expression of subsets of M genes between Y and A/N subtypes (**Figure 1E**), and that each M path involves upregulation of a distinct set of M genes. Further analysis with the two leading M PCs showed that M markers such as VIM contributed to PC1 (M1) significantly, whereas M2 was supported by other M markers such as CXCR4 (**Figure 1F**). While both subsets of M genes are well recognized M genes (Cheng, et al., 2018; Mendez, et al., 2010), they showed distinct patterns across different SCLC cells.

We next use example scRNA-seq datasets from two separate time-course experiments, each profiling EMT induction in A549 cells treated TGF-β (12,911 cells) (Cook and Vanderhyden, 2020). With our E scores and M scores and GMM-based clustering in E-M space, we found that cells at the hybrid EMT state are mainly distributed at the transition time points of the experiments (**Figure 1G and H**), and that the progression of cell states over time was strongly correlated with the downregulation of E genes and upregulation of M genes with the treatment of TGF-β (**Figure 1H**). This trend was also consistent with the progression of M scores (M1 from nnPCA) over pseudotime (**Figure 1I**, blue), which is more widely available from scRNA-seq datasets compared to true time labels. Interestingly, independent analysis of A549 cells treated with EGF (12,435 cells) from the same study did not show such a correlation (**Figure 1I**). This observation is consistent between nnPCA and other methods (**Figure S1**). Finally, we identified several gene sets whose scores were strongly positively/negatively correlated with M (i.e. EMT) scores (**Figure 1J** and **Figure S2**).

## 4. Conclusions

EMTscore provides flexible, versatile and easy-to-use functionality that allows in-depth analyses with gene set scores for EMT progression and publication-quality visualizations. The package is suitable for both bulk and sing-cell omics datasets. A default nnPCA-based method enables identification of multiple, divergent EMT pathways that are supported by different subsets of M genes, and this method is efficient for screening pathways that are associated with EMT.

## Author contributions

Haimei Wen: Methodology, Software, Writing—original draft, Writing—review & editing. Leonidas Bleris: Methodology, Writing—review & editing. Zhao Sun: Methodology, Writing—review & editing, Tian Hong: Project administration, Supervision, Resources, Methodology, Writing—original draft, Writing—review & editing.

