## Supplementary information for "EMTscore infers divergent EMT pathways from omics data and enables rapid screening for correlated gene sets"

**Supporting text for gene set scores based on nonnegative PCA**

To quantify the divergent progression of EMT, we performed nonnegative PCA (nnPCA) using the gene sets mentioned above (Panchy, et al., 2021; Panchy, et al., 2022). nnPCA determines the approximately orthogonal axes with non-negative coefficients (loadings) for features (genes). Variances of projections of data points (cell or cell lines) onto these axes are maximized via an optimization method (Sigg and Buhmann, 2008). In brief, the vectors of weights, $w$, used to define the first principal component of PCA is defined such that it maximizes the variance of the first component, i.e:

$${arg max}_{\boldsymbol{w}} w^{T}Cw (1)$$

where $C$ is the covariance matrix of the original data set $X$ and $w$ is unit vector (${||w||}^{2} = 1$)). In our case, $X$ is an $m$ by $n$ matrix of expression values where $m$ is the number of samples and $n$is the number of genes in the selected gene set. This method for determining $w$ can be treated as an expectation maximization problem where the original data is projected using the current estimate of $w$ ($y = Xw_{t}$) and this projection is used to re-estimate $w$ using the following minimization step:

$$w_{t+1}={arg min}_{w}\sum_{n=1}^{N} ||x_{n}-y_{n}w{||}_{2}^{2} (2)$$

where $x_{n}$ are the rows of the original data and $y_{n}$ are the rows of the projected data (Sigg and Buhmann, 2008). This expectation-maximization formulation allows additional constraints on $w$, including forcing the component values to be non-negative. Note that the non-negativity constraint applies only to the weight components such that negative scores can still exist if there are negative values in underlying data, such as those produced by centering expression data to zero which we did for all nnPCA inputs. Subsequent components are calculated in the same way, under the constraint that they are orthogonal to the preceding ones.

To select the principal components (PCs) that best represent the EMT programs across SCLC subtypes, we selected the top two PCs that have the highest variances explained for individual samples (cells or cell lines). To identify genes that contribute significantly to each PC, we used their nonnegative loading values in the rotation matrix from nnPCA and used the top ranking genes for each PC.

**Supplementary Figures**

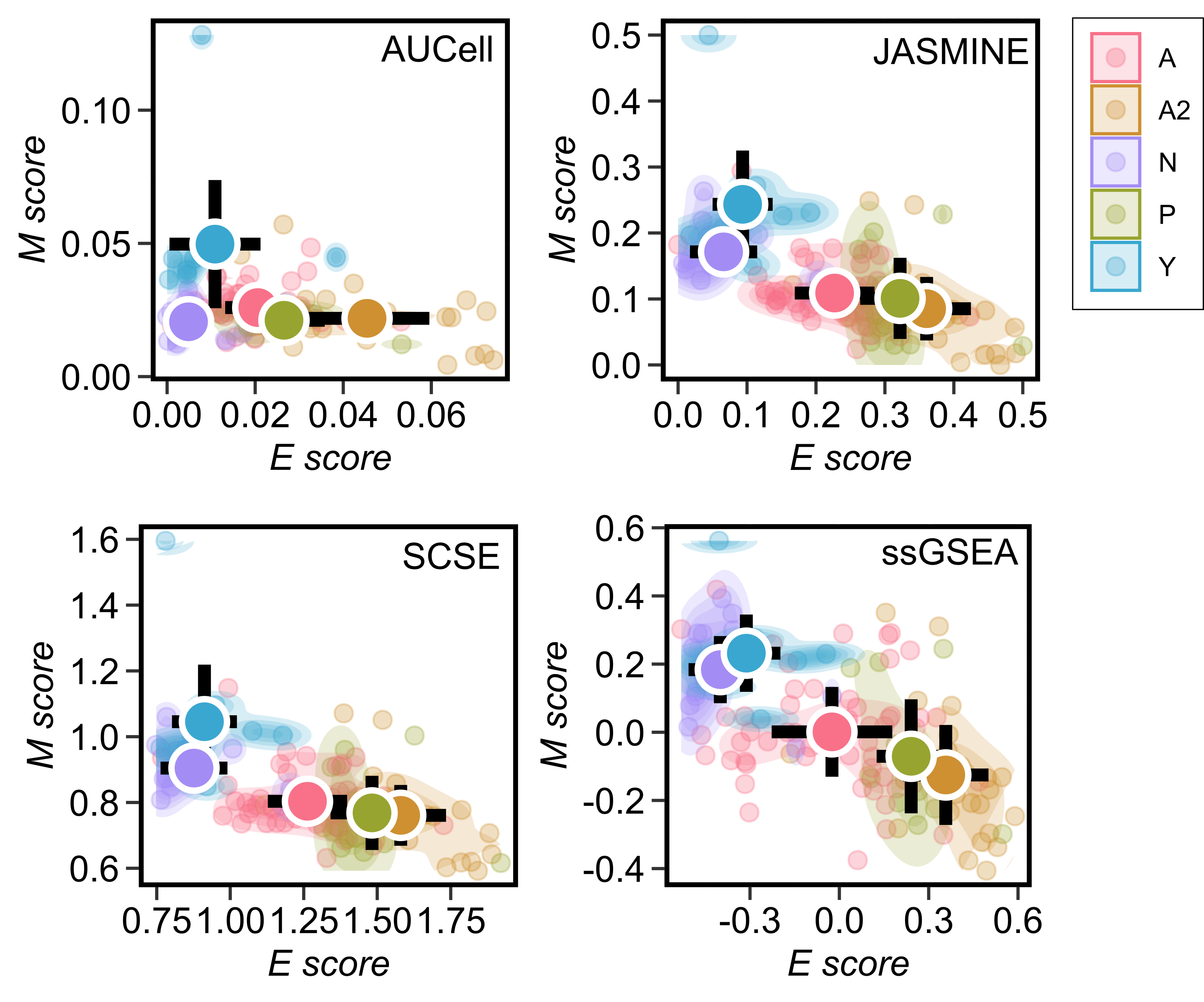

**Figure S1**. **E and M scores computed by four alternative methods**. Bulk RNA-seq data for 120 SCLC cell lines with annotated corresponding tumor types and Panchy et al. M and E gene sets were used (Groves, et al., 2022; Panchy, et al., 2022). Four alternative gene set scoring methods were implemented in EMTscore (Aibar, et al., 2017; Barbie, et al., 2009; Noureen, et al., 2022; Pont, et al., 2019).

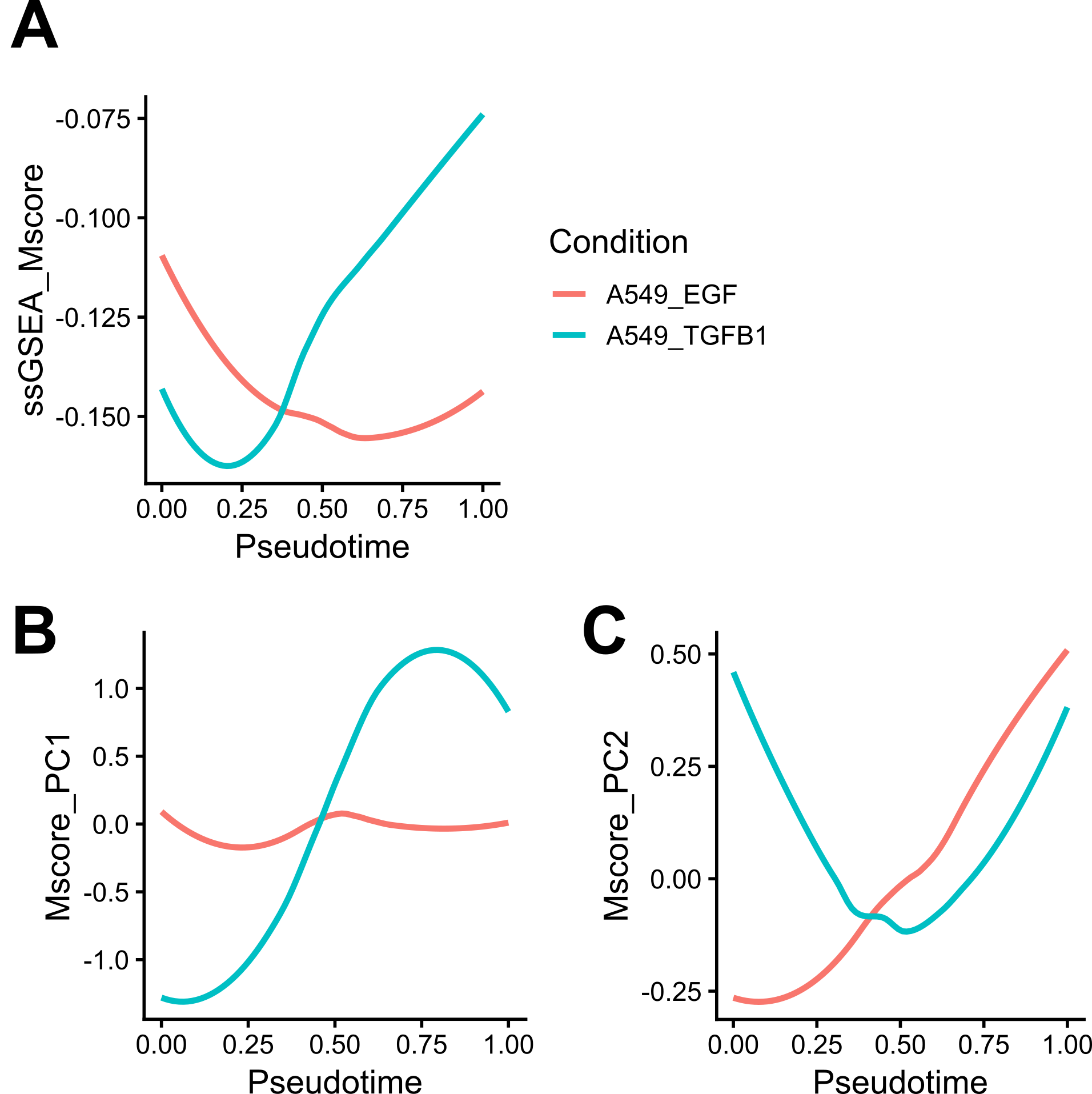

**Figure S2**. **Divergence of EMT scores with respect to pseudotime**. Single-cell RNA-seq (scRNA-seq) data for TGF-β treated A549 cells and EGF treated A549 cells were analyzed with EMTscore separately. ssGSEA (single M score. **A**) and nnPCA (**B**: M1 and **C**: M2) were used to calculate M scores. Panchy et al. M gene set was used (Panchy, et al., 2022). These EMT scores are plotted as functions of pseudotime.

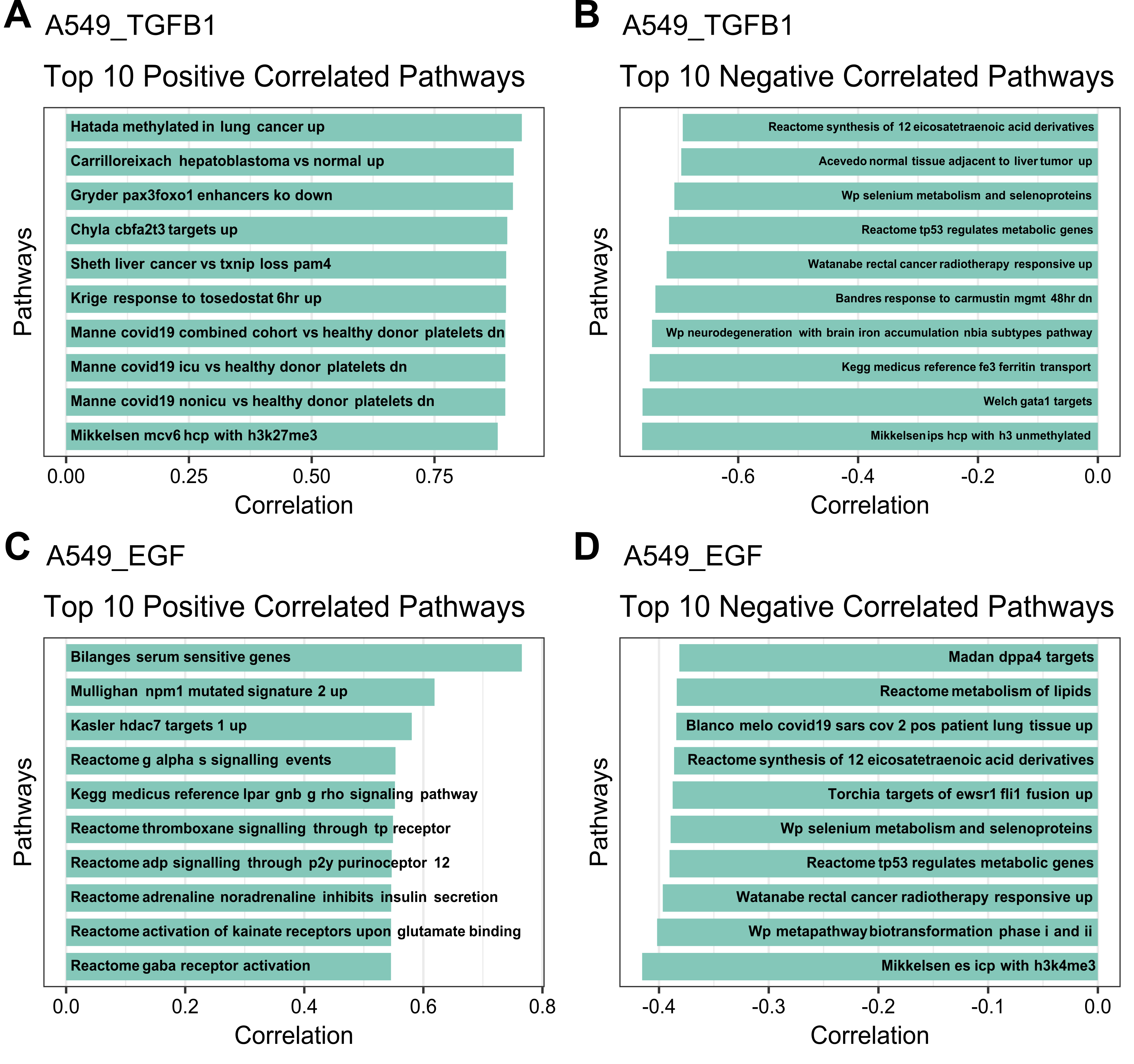

**Figure S3**. **Top gene sets correlated with M scores**. Single-cell RNA-seq (scRNA-seq) data for TGF-β treated A549 cells and EGF treated A549 cells were analyzed with EMTscore separately. M1 scores for TGF-β treatment and M2 scores for EGF treatment were used to screen MsigDB C2 gene sets whose scores are correlated with EMT. Panchy et al. M gene set was used (Panchy, et al., 2022). Gene sets with more than 30% overlapped genes with the M gene set were filtered out for screening.
